# Novel bacterial lineages associated with boreal moss species

**DOI:** 10.1101/219659

**Authors:** Hannah Holland-Moritz, Julia Stuart, Lily R. Lewis, Samantha Miller, Michelle C. Mack, Stuart F. McDaniel, Noah Fierer

## Abstract

Mosses are critical components of boreal ecosystems where they typically account for a large proportion of net primary productivity and harbor diverse bacterial communities that can be the major source of biologically-fixed nitrogen in these ecosystems. Despite their ecological importance, we have limited understanding of how microbial communities vary across boreal moss species and the extent to which local environmental conditions may influence the composition of these bacterial communities. We used marker gene sequencing to analyze bacterial communities associated with eight boreal moss species collected near Fairbanks, AK USA. We found that host identity was more important than site in determining bacterial community composition and that mosses harbor diverse lineages of potential N^2^- fixers as well as an abundance of novel taxa assigned to understudied bacterial phyla (including candidate phylum WPS-2). We performed shotgun metagenomic sequencing to assemble genomes from the WPS-2 candidate phylum and found that these moss-associated bacteria are likely anoxygenic phototrophs capable of carbon fixation via RuBisCo with an ability to utilize byproducts of photorespiration from hosts via a glyoxylate shunt. These results give new insights into the metabolic capabilities of understudied bacterial lineages that associate with mosses and the importance of plant hosts in shaping their microbiomes.

## Introduction

Mosses, like their cousins, the vascular plants, associate with a broad diversity of microbes, including, bacteria, fungi, and other microbial eukaryotes (Lindo and Gonzalez, 2010). These moss- microbe associations are particularly relevant to terrestrial nitrogen (N) and carbon (C) cycling in northern ecosystems, where mosses are ubiquitously distributed and can be responsible for as much as 50% of ecosystem net primary productivity (Turetsky *et al.*, 2012). Moss-associated N^2^-fixing bacteria are often the primary source of ecosystem N inputs in boreal forests (DeLuca *et al.*, 2002) and moss-associated microbes can have important influences on ecosystem C dynamics via methane oxidation, especially in peatlands, one of the largest natural sources of atmospheric methane (Kip *et al.*, 2010). Together, the importance of moss-microbial communities to terrestrial biogeochemistry and unique features of bryophyte biology make boreal moss communities a useful system for investigating the interactions between host species identity, microbial community structure, and ecosystem function.

Mosses are ubiquitous across boreal forests, with distributions spanning ecologically important environmental gradients (Lindo and Gonzalez, 2010; Turetsky *et al.*, 2012). Moss diversity in these forests can be quite high (Geffert *et al.*, 2013), and different moss species often grow interspersed at a given location, creating many abundant and naturally occurring ‘common-garden’ experiments for testing how host identity and local environmental conditions influence the assembly of moss-associated microbial communities. Furthermore, mosses, unlike more commonly-studied vascular plants, have simpler leaf-tissue structures and do not possess roots thus representing a comparatively homogeneous host environment, reducing the need to control for inter-tissue spatial variation (a problem when studying vascular plants, e.g. Leff *et al.*, 2015). Mosses are also small enough that an entire plant can be sampled for microbial analyses, an impossible task for most larger plants. Together these traits make mosses a useful study system for investigating the impacts of environmental and host factors on microbial community structure and the contributions of these moss-associated microbial communities to ecosystem function.

Despite their potential biogeochemical importance and their utility as a model system, we still know surprisingly little about the structure and function of those microbial communities associated with boreal mosses. Much of the previous research has focused on a handful of abundant boreal moss species, in particular, members of the peat-moss genus *Sphagnum,* and the dominant feather mosses, *Pleurozium schreberi* and *Hylocomium splendens* (e.g. DeLuca *et al.*, 2002; Opelt *et al.*, 2007; Zackrisson *et al.*, 2009; Ininbergs *et al.*, 2011; Bragina, Berg, *et al.*, 2012; Bragina *et al.*, 2015). While these species are among the most abundant mosses in many boreal forests, the microbes living in association with less abundant hosts may be equally as important contributors to key ecosystem processes (Rousk *et al.*, 2015). Previous work on the abundant species makes it clear that boreal mosses possess surprisingly diverse microbial communities, containing not only well-studied *Cyanobacteria* but also novel and undescribed lineages within the *Alphaproteobacteria* sub-phylum and the *Verrucomicrobia* phylum among others (Bragina *et al.*, 2015). However, it is unclear whether local environmental factors govern the microbial community assembly, or whether a moss species hosts a characteristic microbial community, regardless of its abiotic environment.

It is well-known that mosses can harbor N^2^-fixing bacteria and N^2^-fixation rates of *Sphagnum* and the feather mosses, *Pleurozium schreberi* and *Hylocomium splendens,* can be quite variable across season (DeLuca *et al.*, 2002), forest type (Zackrisson *et al.*, 2004), and moss species (Leppänen *et al.*, 2015) with reported rates ranging from 0.3-4 kg N ha^-1^ yr ^-1^ (DeLuca *et al.*, 2007). A number of previous studies have focused on selected moss-associated N^2^-fixing bacterial taxa, particularly taxa within the *Cyanobacteria* phylum (reviewed in Rousk, Jones, *et al.*, 2013), and how the composition or diversity of these cyanobacteria relate to N^2^-fixation rates (Ininbergs *et al.*, 2011). However, *Cyanobacteria* are unlikely to be the only moss-associated bacteria capable of N_2_-fixation. For example, Bragina *et al.* (2012) found that nitrogenase sequence libraries from bacteria on two *Sphagnum* species were dominated by alphaproteobacterial sequences, indicating that non- cyanobacterial N_2_-fixers may play a more important role than previously recognized. In short, it remains unclear which bacteria are responsible for N_2_-fixation in boreal mosses and how the abundances of these N_2_-fixing taxa may vary across different moss species.

Here we characterized the bacterial communities associated with eight common boreal moss species at three different sites using marker gene (16S rRNA gene) amplicon sequencing. We used this dataset to address the following questions: 1) How do bacterial communities vary across different moss species and different sites? and 2) Which microbial taxa from these communities are potential N_2_-fixers? Based on our observation that many of the taxa found in association with the moss species were representatives of bacterial lineages for which little is known (including members of the candidate phylum WPS-2), we then assembled genomes from shotgun metagenomic data to determine the functional attributes of these abundant, ubiquitous, and previously undescribed members of moss- associated microbial communities.

## Experimental Procedures

### Sample Collection

To characterize and compare the bacterial communities associated with different moss host species and their potential contributions to N_2_-fixation, we collected samples of eight boreal moss species (the closely related pleurocarpous mosses *Pleurozium schreberi* (Brid.) Mitt., *Hylocomnium splendens* (Hedw.) Shimp, *Sanionia uncinata* (Hedw.) Loeske, and *Tomenthypnum nitens* (Hedw.) Loeske, and the successively more distantly related species, *Aulacomnium palustre* (Hedw.) Schwägr. and *A. turgidum* (Wahlenb.) Schwägr., *Dicranum elongatum* Schleich. ex Schwägr., and *Sphagnum capillifolium* (Ehrh.) Hedw.). The samples were collected from three black spruce (*Picea mariana*)-dominated sites that were each at least a kilometer apart within the arboretum of the University of Alaska Fairbanks during late July 2014. The selected moss species are both common in boreal forests and represent lineages spanning the phylogenetic diversity of mosses. At each of the three sites we collected one sample for each of the eight species (except *Aulacomnium turgidum* and *Dicranum elongatum*, which could only be found at two sites), resulting in a total of 22 samples for analysis. Each sample was divided into four subsamples of ten stems each for microbial community analysis, isotopic enrichment, natural abundance voucher, and taxonomic voucher specimens (See Table S1). For each species within each site, a clump of ramets were removed from a monospecific patch and carefully sorted and cleaned with a gloved hand. Brown or decaying material was removed from the bottom so that each ramet was approximately 5 cm in length and included the apical meristem. Each sample was then divided into four subsamples of ten stems each for microbial community analysis, isotopic enrichment, natural abundance voucher, and taxonomic voucher specimens (See Table S1). Samples collected for microbial analyses were placed in a cooler on blue ice in the field and returned to the lab where they were frozen (-20°C) within 2 h of collection. Samples for the N_2_-fixation rate assays were also placed in the coolers until the measurements were started, usually within 2 h of sample collection. Natural abundance voucher specimens were returned to the lab and dried at 60°C for 48 h within 2 h of collection.

### N_2_-Fixation Measurements

To quantify field rates of N_2_-fixation for each sample, we used an isotopic enrichment approach modified from Ruess *et al.* (2009, 2013). Each sample was placed in a 60 mL translucent polycarbonate syringe that was depressed to contain 10 ml of air. We then added 10 mL of ^15^N_2_ (98% enriched, Cambridge Isotope Laboratories, Inc., U.S.A.) and sealed the syringe with a stopcock. Sealed syringes from all sites were placed in site A, where they were incubated for 24 h under similar light and temperature conditions. After 24 h, mosses were removed from the syringes and placed in a 60 ^o^C oven for 48 h. Both enriched incubation samples and natural abundance samples were ground to a homogenous powder. Nitrogen and carbon concentrations and atom% ^15^N and ^13^C values for both enriched incubation samples and natural abundance voucher specimens were measured on a Costech ECS4010 coupled to a Thermo Scientific Delta V Advantage Isotope ratio mass spectrometer in Mack’s lab at Northern Arizona University. Nitrogen fixation was calculated by comparing the ^15^N values from enriched and control samples (see Jean *et al.* (2017) for details).

### Amplicon-Based Bacterial Community Analysis

To analyze the bacterial communities associated with each of the collected moss specimens, we PCR amplified and sequenced a portion of the bacterial and archaeal 16S rRNA marker gene. First, to minimize any bias from micro-spatial differences along the moss tissue, we homogenized each sample (ten moss stems per sample, approximately 0.25 g of tissue) with liquid N_2_ under sterile conditions. We then extracted DNA from each homogenized moss sample using the MoBio Power Soil DNA extraction kit (MoBio Laboratories, Carlsbad, CA). After extracting DNA, we used the 515f / 806r primers to PCR amplify the V4-V5 region of the 16S rRNA gene (Caporaso *et al.*, 2012). For each sample, we used a unique primer pair that included a 12-bp barcode and Illumina sequencing adapters to allow for multiplexed sequencing. To minimize amplification of mitochondrial and chloroplast DNA, we used PNA (peptide nucleic acid) clamps during PCR amplification (Lundberg *et al.*, 2013). During both DNA extraction and PCR amplification, we included negative controls to check for potential contaminants introduced during those steps. We prepared the samples for sequencing by normalizing the concentrations of PCR products across all samples using the ThermoFisher Scientific SequalPrep Normalization plate (Thermo Fisher Scientific Inc. USA) and pooled the amplicons together. We sequenced all samples on the Illumina MiSeq platform running the 2 x 150 bp paired-end chemistry at the University of Colorado Next Generation Sequencing Facility. The *Hylocomium splendens* amplicon sequences were sequenced on a separate run. Because we cannot control for run-to-run variation we have excluded these samples from downstream statistical analyses. A cursory analysis indicates that the *Hylocomnium* microbiome is statistically distinguishable from the other mosses in our sample, however the community was not marked by novel species relative to the other samples we analyzed.

We used the approach described by Leff *et al.* (2015) to analyze the 16S rRNA sequence data. Briefly, we removed adapters from the raw reads using cutadapt (paired, -O 1) (Martin, 2011) and demultiplexed the reads using a custom in-house python script (“prep_fastq_for_uparse_paired.py” at: https://github.com/leffj/helper-code-for-uparse/). Then, using USEARCH v.8 (Edgar, 2010), we merged, quality filtered (“maxee rate” = 0.005) and dereplicated the reads to create a fasta file containing each unique amplicon sequence. We also removed singletons (sequences found only once across the entire sample set) from the dereplicated reads. We created a de novo database from our sequences using the UPARSE pipeline (Edgar, 2013) (implemented with USEARCH v.8) by clustering reads at ≥97% sequence similarity level (which we refer to as ‘phylotypes’). As a quality-control measure, we removed representative sequences from phylotypes that were less than 75% similar to any sequence in the Greengenes database (version August 2013) (McDonald *et al.*, 2012). We assumed these highly divergent sequences to be chimeric, a product of non-specific amplification, or of insufficient quality.

To generate phylotype counts, we mapped the merged reads back to the *de novo* database (command “-usearch_global”, -id 0.97) and used a custom python script (“create_otu_table_from_uc_file.py” at: https://github.com/leffj/helper-code-for-uparse/) to generate a table of phylotype counts per sample from the USEARCH mapping output. We classified the reads with the RDP Naïve Bayesian classifier (Wang *et al.*, 2007) against the Greengenes database (McDonald *et al.*, 2012) and removed any residual chloroplast and mitochondrial sequences. To control for differences in read depth (sequencing “effort”) across samples, we randomly selected 6000 bacterial and archaeal 16S rRNA gene reads per sample prior to downstream analyses. The 6000 read cut-off was chosen based on the read depth of the sample with fewest reads after mitochondria and chloroplast removal. The filtered reads can be found on FigShare (DOI: https://doi.org/10.6084/m9.figshare.5594527).

To measure differences between communities across host species and sites, we calculated pairwise Bray-Curtis dissimilarities on the square-root transformed phylotype table and tested for differences between species and sites using a PERMANOVA test. All statistical analyses were conducted in R (R Core Team, 2017, package “Vegan”, R Core Team, 2017). We used metaMDS (R package “Vegan”) to generate NMDS plots from the dissimilarity matrix.

### Identifying Nearest Isolated Representatives and Classifying Putative N2-fixing Bacteria

To identify the nearest isolated representatives of the 30 most abundant bacterial phylotypes, we used the RDP (Ribosomal Database Project) SeqMatch tool which finds the nearest relatives by comparing the percent of shared sub-sequences between a query and the RDP isolate database (Cole *et al.*, 2014). We then downloaded the nearest-neighbor representative sequences from RDP and used MUSCLE (Edgar, 2004) to create an alignment of the isolate sequences and representative sequences from the 30 most abundant bacterial phylotypes, using FastTree (Price *et al.*, 2010) to generate the final tree. We defined “putative N_2_-fixing phylotypes” as those phylotypes whose nearest relative (≥97% similarity in the 16S rRNA gene region) has been found to be capable of N_2_-fixation when cultivated in isolation under laboratory conditions (four phylotypes). Alternatively, if the nearest neighbor was < 97% similar but came from a lineage which included only previously-described representative strains known to be capable of N_2_-fixation, the phylotype was also defined as a “putative N_2_-fixer” (two phylotypes).

### Shotgun Metagenomic Sequencing and Analysis

Based on the preponderance of bacterial lineages identified from the 16S rRNA amplicon analyses that came from poorly described and novel bacterial lineages (see below), we generated shotgun metagenomic libraries from the samples of 3 moss species *Aulacomnium turgidum* (2 samples), *Pleurozium schreberi* (3 samples), and *Tomenthypnum nitens* (3 samples). We prepared the metagenomic libraries following the method described in Baym *et al.* (2015). All 8 libraries were then sequenced on the Illumina NextSeq platform running the 2 x 150 bp chemistry at the University of Colorado Next Generation Sequencing Facility. Each sample had an average of 58 million reads. After sequencing, we filtered the raw reads with Sickle (-q 20 -l 50) (Joshi and Fass, 2011) and used Metaxa2 (Bengtsson-Palme *et al.*, 2015) on the filtered reads to verify that the 16S rRNA amplicon data was consistent with the taxonomic composition of the bacterial communities as inferred from the metagenomic data.

To assemble near complete genomes from the shotgun metagenomic data, we used the metagenomic *de novo* assembler, MEGAHIT (Li *et al.*, 2014) (default settings: --min-count 2 --k-min 21 --k-max 99 --k-step 20) to co-assemble all the filtered reads without regard to sample origin (pooled-assembly). Then we sorted the assembled contigs into bins each representing a preliminary genome using the software MaxBin (Wu *et al.*, 2014) (-min_contig_length 1000 -max_iteration 50 -prob_threshold 0.9 -markerset 40). Since MaxBin can take advantage of differences in abundance of identical contigs across samples, we were able to leverage differential abundances of reads from individual bacterial taxa across the pooled assembly to identify contigs belonging to the same organisms, thus facilitating the binning process. After binning, we used Bowtie2 (Langmead and Salzberg, 2012) to map the filtered reads back to each bin. We used CheckM (Parks *et al.*, 2015) to verify the completeness and contamination of each bin (Table S2). Briefly, CheckM measures metagenomic completeness and contamination on the basis of presence and number of conserved single-copy marker genes (genes which typically are present in bacterial genomes only once). The number of marker genes present compared to the number of expected marker genes for a particular bacterial lineage is used as a measure of completeness while the number of copies of a marker gene indicates contamination. We considered bins that were greater than 70% complete and less than 10% contaminated to be “high-quality” bins (as per the top two levels of quality identified in Parks *et al.* (2015).

Because few of our high-quality bins contained assembled 16S rRNA sequences (a common problem, see Miller *et al.*, 2011), we used three separate methods to confirm the taxonomic identity of bins. First, we used the program EMIRGE (Miller *et al.*, 2011) to assemble full-length 16S rRNA sequences from the raw metagenomic reads and used USEARCH (Edgar, 2010) to match contigs from the bins to the EMIRGE-assembled full-length 16S rRNA sequences. Second, to verify this result with a technique that did not rely on genome assembly from metagenomic data, we used Metaxa2 (Bengtsson-Palme *et al.*, 2015) to independently identify fragments of the 16S rRNA gene in the binned contigs and classified those fragments against the Greengenes database using USEARCH (Edgar, 2010). Finally, we used the automatically-generated concatenated marker gene phylogeny available from CheckM to confirm that bins were clustered on a tree in the appropriate clade as indicated by analysis of their 16S rRNA gene sequences. After identifying the organisms represented by the bins, we used the Joint Genome Institute’s Integrated Microbial Genomes (IMG) analysis pipeline (Markowitz *et al.*, 2014) to identify and assign functions to the genes in the assembled bins. The assembled metagenomic reads are available through the IMG portal (IMG submission ID 115847).

## Results and Discussion

### Moss Species Identity Drives Microbiome Composition

Our analyses of moss-associated bacterial 16S rRNA amplicons show that mosses host diverse and species-specific bacterial communities. After quality filtering, but prior to chloroplast and mitochondrial filtering, we obtained 7947 to 16590 16S rRNA gene reads per sample with an average sequencing depth of 11430 reads. The chloroplast and mitochondrial sequences that were removed made up only a small percentage of the reads (10% and 5%, respectively). Samples had an average richness of 924 phylotypes per sample and were dominated by eight bacterial phyla. Archaeal sequences were extremely rare and made up less than 0.001% of all reads. We found similar archaeal abundances in the primer-independent metagenomic data suggesting that the low archaeal abundance in the amplicon data was not a product of primer biases. The dominant bacterial phyla, as measured by average read abundance across all samples, were *Proteobacteria* (44.8% of reads across all samples), *Acidobacteria* (10.8%), *Verrucomicrobia* (9.8%), *Bacteroidetes* (9.3%), *Cyanobacteria* (6.5%), Candidate phyla WPS-2 (5.7%), *Planctomycetes* (5.2%), and *Actinobacteria* (4.2%) (Figure 1). The most abundant phylotypes identified across the entire sample set included those assigned to the *Acetobacteraceae* (9.5%), *Acidobacteriaceae* (8.2%), *Sinobacteraceae* (8.2%) and *Nostocaceae* (5.2%) families as well as many phylotypes that could only be classified to the phylum or class level of resolution (including those in the WPS-2, and *Verrucomicrobia* phyla, Figure 1). These phyla and families have been found in other microbial studies of mosses; notably *Acetobacteraceae* and *Acidobacteriaceae* were found to be abundant in sphagnum mosses (Bragina, Berg, *et al.*, 2012) and *Nostocaceae* have long been studied in association with feather mosses(DeLuca *et al.*, 2002) and *Sphagnum* (Bay *et al.,* 2013; Kostka *et al.,* 2016). Members of the WPS-2 and *Verrucomicrobia* phyla have previously been found in association with sphagnum mosses in bogs (Bragina *et al.*, 2015).

**Figure 1:**
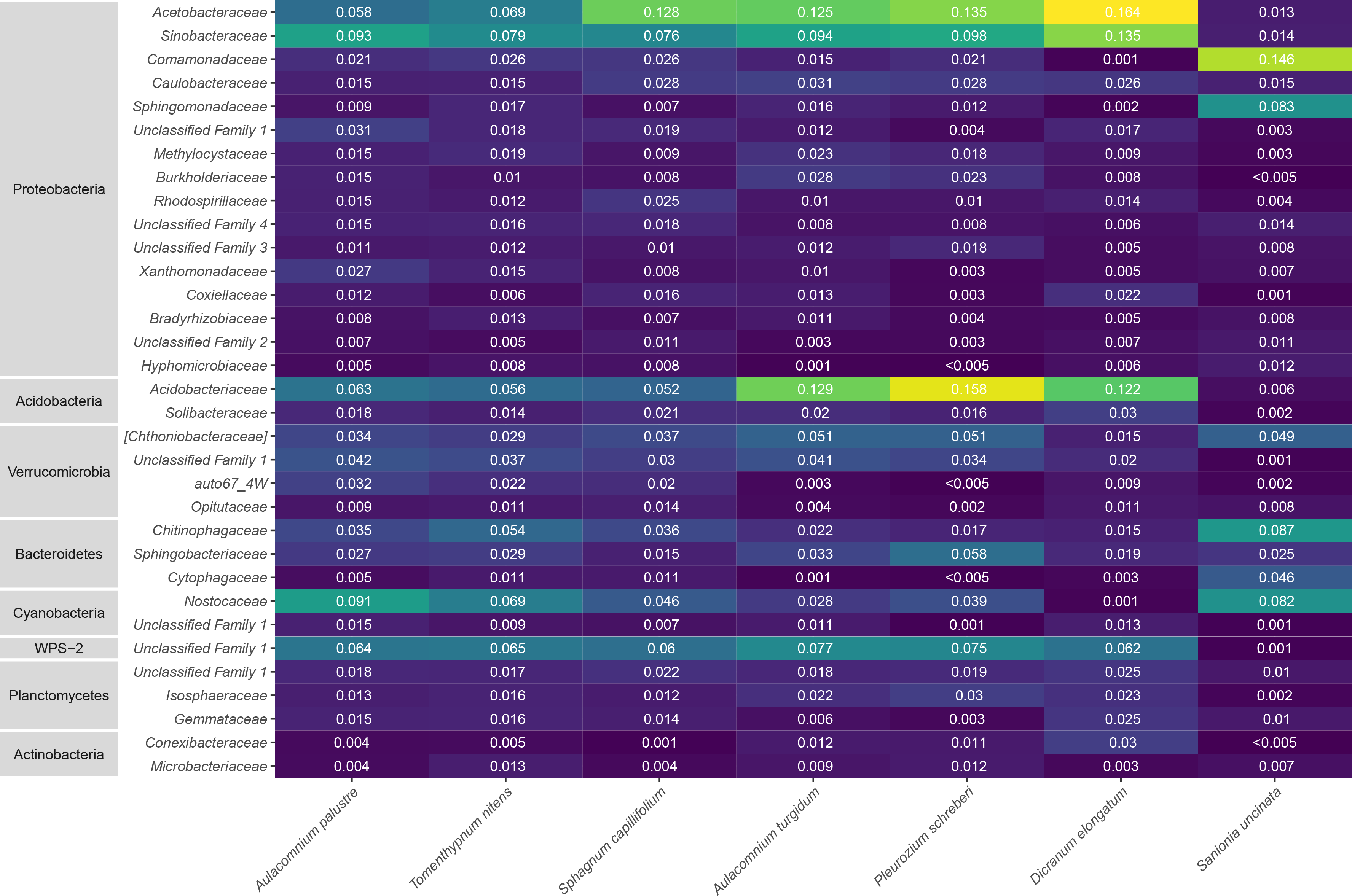
Heat map showing the relative abundances of the most abundant bacterial families across the 7 moss species. Phylum-level classifications for each family are noted on the left. Higher relative abundances are indicated in brighter colors. Families with no official classification are noted as “Unclassified Family.” Host moss species are indicated along the x-axis and ordered according to overall bacterial community similarity.

Moss species identity was more important than site in determining the composition of the moss-associated bacterial communities. Across all samples, moss species contributed to 63% of the variation in community composition (Permanova R^2^ = 0.63, p < 0.001), while site was not a significant source of variation (Permanova R^2^ = 0.076, p = 0.21) (Figure 2). This finding is consistent with studies in vascular plants which find that species identity is generally more important than site in shaping phyllosphere community composition (Redford *et al.*, 2010; Laforest-Lapointe *et al.*, 2016). However, sites in this study were situated within a relatively small area. It is possible that across larger scales, differences in local environmental conditions may have a stronger influence on the observed bacterial communities. Across all moss species studied here, *Sanionia uncinata* harbored particularly distinct bacterial communities. More than half (59%) of phylotypes found on *Sanionia uncinata* were not found in any of the other moss species and, of the 30 most abundant phylotypes highlighted in Table S3, only 77% were present on *S. uncinata* compared to more than 90% found across every other species (Figure 3). While most of the mosses were dominated by *Acetobacteraceae* (*Alphaproteobacteria*), *Acidobacteriaceae* (*Acidobacteria*), and *Methylacidiphilales* (*Verrucomicrobia*), the moss species *Sanionia uncinata* had a low abundance or complete absence of these taxa. Instead, *Sanionia uncinata* was dominated by *Comamonadaceae* (*Betaproteobacteria*), *Nostocacaceae* (*Cyanobacteria*) and *Chitinophagageae* (*Bacteroidetes*). The distinctiveness of the *S. uncinata* microbiome does not appear to reflect any aspect of phylogenetic relatedness among the sampled moss species. For instance, the pleurocarpous *S. uncinata* has much closer phylogenetic affinities with *P. schreberi* and *T. nitens*, yet the phylogenetically distant *Sphangnum capillifolium* hosts a microbiome much more similar to *P. schreberi* and *T. nitens* than does *S. uncinata*. The factors driving these differences between *S. uncinata* and the other moss species remain uncertain.

**Figure 2:**
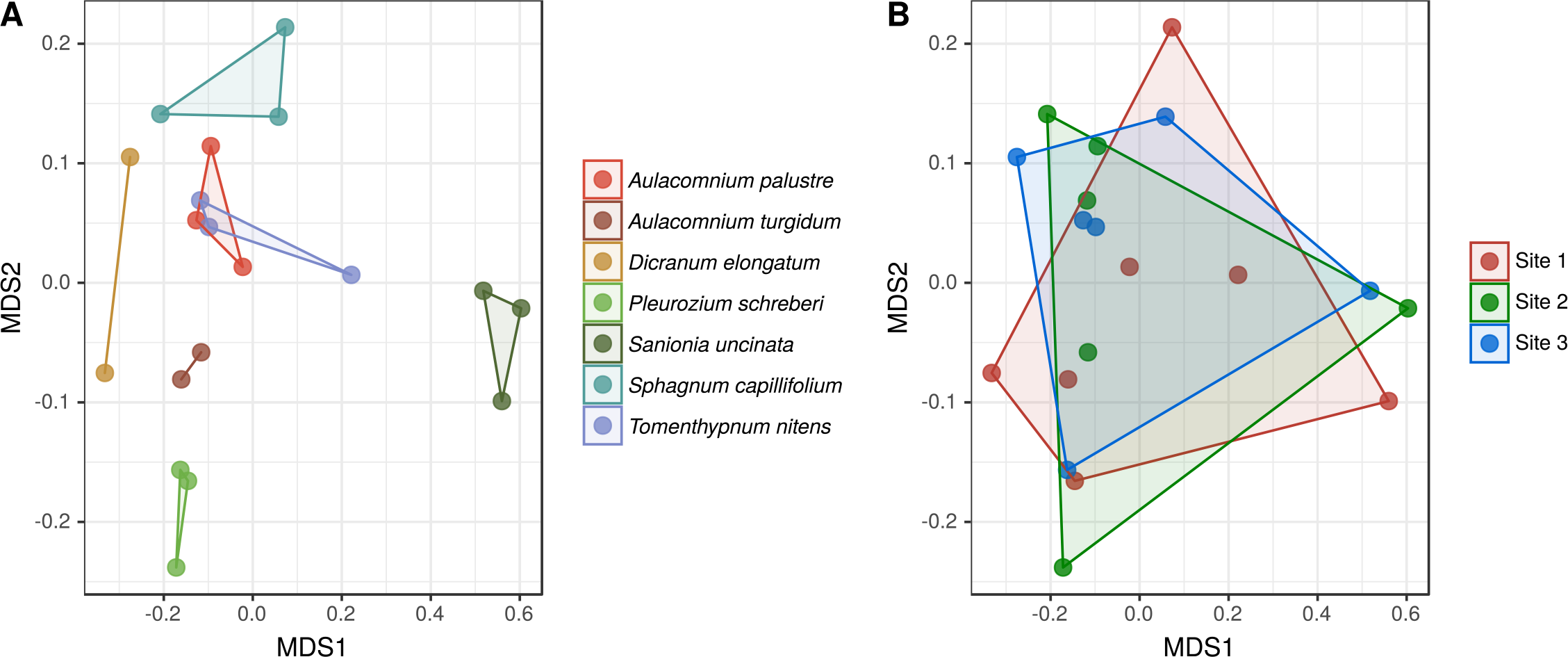
Non-metric multidimensional scaling (NMDS) plot of bacterial community dissimilarities across samples (measured using the Bray-Curtis distance metric). Samples from the same host species tend to cluster together (A), while those from the same site were spread across ordination space (B). A PERMANOVA test revealed that species identity contributes to 63% of the variation in community composition (R^2^ = 0.63, p < 0.001), while site is not a significant source of variation between these communities (R^2^ = 0.08, p = 0.21).

### Evidence for multiple groups of N2-fixing bacteria in boreal mosses

Using stable-isotope enrichment, we measured N_2_-fixation rates and found that all of the moss species were actively fixing N_2_ (Figure S1), though rates varied by a factor of 100 across samples (from 0.98 to 100 μg N • g dry weight moss^-1^ day^-1^). Importantly, we find that many moss microbiomes which were assumed to be non-N^2^-fixing (the genus *Dicranum*, for example, Gundale *et al.* (2011) do, in fact, appear to associate with bacteria capable of fixing N_2_ at rates comparable to those of feather mosses, which are well-known to host bacteria capable of sustaining relatively high rates of N_2_-fixation (DeLuca *et al.*, 2002; Turetsky *et al.*, 2012). Consistent with other studies, *Pleurozium schreberi* and *Sanionia uncinata* had the highest average rates of N_2_-fixation at 46.1 and 52.4 μg N • g dry weight moss^-1^ day^-1^, respectively. Our measured rates of fixation are within the expected range previously reported for *Sphagnum* mosses and *Pleurozium schreberi* (Vile *et al.*, 2014). However, because we used a ∂^15^N method as opposed to an acetylene reductase assay, and because of the lack of reported N_2_ fixation studies for some of the moss species in our study, it is difficult to directly compare all of our measurements of N_2_-fixation rates to those reported previously (e.g. DeLuca *et al.*, 2002; Zackrisson *et al.*, 2004; Gundale *et al.*, 2012).

Given that all of the sampled moss species had microbial communities with measurable N_2_-fixation activities, we then used the 16S rRNA gene survey data to determine which taxa might be contributing to the N_2_-fixation. We found three cyanobacterial and three alphaproteobacterial lineages that were reasonably abundant across nearly all moss species, with all of these lineages having previously been identified as capable of N_2_-fixation. Specifically, the putative N_2_-fixers in our samples were from the *Nostocaeae*, *Acetobacteraceae*, *Bradyrhizobiaceae*, and *Methylocystaceae* families (Figure 3). Previous studies have focused on the role of *Cyanobacteria* in bryophyte-associated N_2_- fixation, but one study in *Sphagnum*, found that 50% of bacterial cells colonizing *Sphagnum* were from *Alphaproteobacteria* and that the *nifH* gene libraries for these species were dominated by alphaproteobacterial, rather than cyanobacterial sequences (Bragina, Maier, *et al.*, 2012). Interestingly, these two groups of putative N_2_- fixers showed contrasting patterns of abundance between *Sanionia uncinata* and *Pluerozium schreberi* (the two moss species with the highest rates of N_2_-fixation). These two hosts showed relatively similar average rates of N_2_-fixation (52.4 μg N g^-1^day^-1^ and 46.1 μgN g^-1^day^-1^), yet *Sanionia uncinata* was dominated by *Cyanobacteria* (79%) and showed low relative abundances of the alphaproteobacterial N_2_-fixers (7.2%), while *Pleurozium schreberi* showed the opposite pattern (6.8% cyanobacterial, 21.4% alphaproteobacterial). Together, these results suggest that multiple bacterial taxa may contribute to the measured N_2_-fixation activity in these boreal moss species and, even though cyanobacterial N_2_-fixers have received the bulk of the attention in previous studies (e.g. Ininbergs *et al.*, 2011; Rousk, Rousk, *et al.*, 2013), they are not the only N_2_-fixing bacteria that associate with boreal mosses. To identify which of these putative N_2_-fixing lineages are responsible for the measured N_2_-fixation activities in these moss species, future work using stable isotope probing-based approaches would be necessary (e.g. Buckley *et al.*, 2007; Jehmlich *et al.*, 2010).

**Figure 3:**
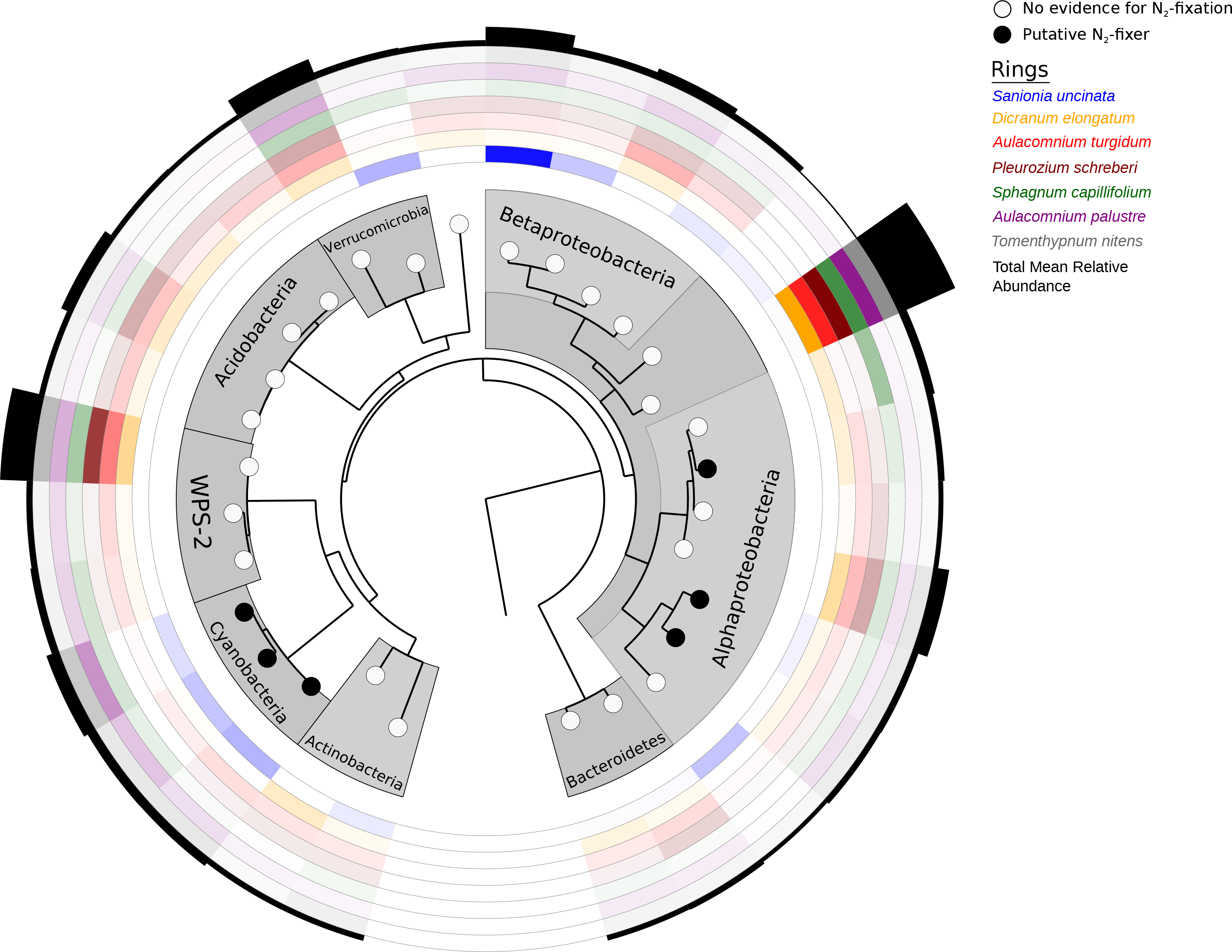
A phylogenetic tree of the top 30 bacterial phylotypes (circles) and their N_2_-fixing potential. Putative N_2_-fixers (black circles) were identified by their relatedness to known N_2_ –fixers (see text). The putative N_2_-fixers observed in our moss samples include representatives from the cyanobacterial and alphaproteobacterial groups. The phylotypes represented here were chosen as they were consistently the most abundant phylotypes across all moss species and together account for 35% of 16S rRNA gene reads in the dataset. Each inner colored ring represents a different species of moss, from the inside ring to the outer ring they are: *Sanionia uncinata* (blue), *Dicranum elongatum* (yellow), *Aulacomnium turgidum* (bright red), *Pleurozium schreberi* (dark red), *Sphagnum capillifolium* (green), *Aulacomnium palustre* (purple), *Tomenthypnum nitens* (gray). The opacity of the colored rings represents the relative abundance of the different phylotypes (log transformed and scaled between 0 and 1) within each moss species. The black outer ring represents the total relative abundance of each phylotype across all host species (also log transformed and scaled between 0 and 1).

### Boreal mosses harbor abundant, undescribed bacterial lineages

The mosses we studied hosted an unexpected abundance of understudied bacterial lineages. Abundant phylotypes for which there were no close matches (<97% 16S rRNA sequence similarity) to previously cultivated and described bacteria are summarized in Table S1. Of these phylotypes, several of the most abundant phylotypes were from the phylum *Verrucomicrobia* and candidate phylum WPS- 2. These phylotypes had no closely related cultivated representatives beyond the phylum-level of resolution, yet they were consistently among the most abundant phylotypes in many of the samples (Table S1, Figure S2). Further analysis of the metagenomic data confirmed the observed high relative abundances of WPS-2 and *Verrucomicrobia* and highlight that the abundances observed with the amplicon-based analyses were not a product of primer biases (Figure S3).

*Verrucomicrobia* are widely distributed in soils (Brewer *et al.*, 2016), acidic geothermal environments (Op den Camp *et al.*, 2009), and have also been found in boreal mosses, particularly in those mosses growing in moist, acidic environments such as bogs and peatlands (Dedysh, 2011; Sharp *et al.*, 2014; Bragina *et al.*, 2015). However, the ecologies of these moss-associated taxa remain poorly known. Among our samples, the most common verrucomicrobial divisions were the orders *Methylacidiphilales* (3.24%) and *Spartobacteria* (3.79%). One phylotype from *Methylacidiphilales*, was the third most abundant phylotype overall. Several earlier studies hypothesized that bacteria from *Methylacidiphilales* contribute to both N_2_-fixation and methanotrophy in peatland environments (Bragina *et al.*, 2015; Ho and Bodelier, 2015). We searched the assembled metagenomic contigs for the methanotrophic marker gene (*pmoA*), and we were unable to find any evidence of the methanotrophic marker gene (*pmoA*) associated with *Methylacidiphilales* in our metagenomic sequence data. It is therefore unlikely that *Methylacidiphilales* in these samples are oxidizing methane unless they are using genes that are not currently recognized in databases as being linked to methane oxidation.

In addition to relatively high abundances of poorly-described verrucomicrobial phylotypes, the mosses were also dominated by a single phylotype from the candidate phylum WPS-2 which represented 3.5% of the 16S rRNA sequences analyzed (Figure 3), making it one of the most abundant phylotypes in six of the seven moss species (with the exception of *Sanionia uncinata*). No members of WPS-2 have ever been cultured, and no published genome exists for any member of this candidate phylum. Given how little we know about this phylum, we can only speculate about the ecology of this group from a few previous studies in which members of this phylum have been detected. Representatives from WPS-2 have previously been found in acidic and cold environments, including alpine bog vegetation (Bragina *et al.*, 2015), mineral deposits from a low-temperature acidic spring (Grasby *et al.*, 2013), and acidic natural gas extraction shale (Trexler *et al.*, 2014). Interestingly, Trexler *et al.* (2014) also observed that WPS-2 seems to co-occur with methanotrophs in aquatic mosses and speculated that WPS-2 may be using derivatives of methanotrophy such as carbon dioxide, formaldehyde, or formate. However, we found only two instances of the methane oxidation marker gene (*pmoA*) in our shotgun metagenomes (out of more than 926 million reads), therefore it seems unlikely that the highly abundant members of the WPS-2 phylum found in our samples associate with methanotrophs.

### Draft genome of WPS-2 recovered from metagenomic data

To learn more about the abundant and poorly studied taxa found in these mosses and to provide further insight into the functional attributes of these bacteria, we chose moss samples with particularly high abundances of diverse taxa of interest (*Methylacidiphilae* and WPS-2) for metagenomic analysis. We assembled 61 genome bins from these moss-associated bacterial communities, seven of which passed contamination and completeness standards (Table S2). Despite the apparent abundance of *Methylacidiphilae*, none of the recovered bins were from this phylum. The inability to recover highly abundant strains from metagenomes is fairly common and can be a product of high strain variation (intraspecific variation) (Miller *et al.*, 2011).

Four phyla were represented in the recovered genome bins: *Acidobacteria* (1 bin), *Proteobacteria* (1 bin, *Alphaproteobacteria*), *Cyanobacteria* (2 bins), and WPS-2 (3 bins). The three bins from WPS-2 ranged from 84 – 89% complete and represent the first genomic data available for members of the candidate phylum WPS-2. We estimate that the full genome sizes for these WPS-2 representatives are 3.88, 3.41, and 4.34 Mbp based on the presence of lineage-specific single-copy marker genes.

Because WPS-2 is one of the most abundant groups across our samples and we know so little about the functional attributes of this group, we used gene annotation to try to reconstruct the potential metabolic attributes of the most complete genome bin obtained for WPS-2. The gene annotations revealed that WPS-2 is likely an anoxygenic phototroph, capable of carbon fixation, and able to metabolize the bi-products of photorespiration making it well-suited to life on the surface of a plant (Figure 4). We identified key genes involved in anoxygenic photosynthesis including those encoding the M and L subunits of anoxygenic photosynthetic reaction centers (*pufM* and *pufL*) as well as the gene for the Y subunit of chlorophyllide a reductase (*bchY*), a universal marker gene for BChl- containing anoxygenic phototrophs (Yutin *et al.*, 2005, 2009). Anoxygenic photosynthesis was first recognized for its importance in marine environments, but recently it has also been recognized as a common trait of phyllosphere bacteria found on the surface of vascular plants (Atamna-Ismaeel *et al.*, 2013). There are three types of anoxygenic phototrophs that contain *pufM*-type reaction centers: purple sulfur bacteria, purple non-sulfur bacteria, and aerobic anoxygenic phototrophs (AAP) (Yurkov and Hughes, 2017). Of these three groups, we hypothesize that WPS-2 is an aerobic anoxygenic phototroph since abundant sulfur sources are expected to be extremely limited in boreal forests and the exposed environment of the moss phyllosphere is unlikely to provide the anaerobic environment necessary to support purple non-sulfur bacteria. If WPS-2 is truly an AAP, it would be one of the few observed lineages to possess RuBisCo. Until recently, all other known AAP were thought incapable of RuBisCo-facilitated carbon fixation (Hughes *et al.*, 2017) however, a recent study (Graham *et al.*, 2017) has found four Alphaproteobacteria in a marine metagenomic dataset that appear to possess the ability to fix carbon via the Calvin-Benson-Bassham cycle.

**Figure 4:**
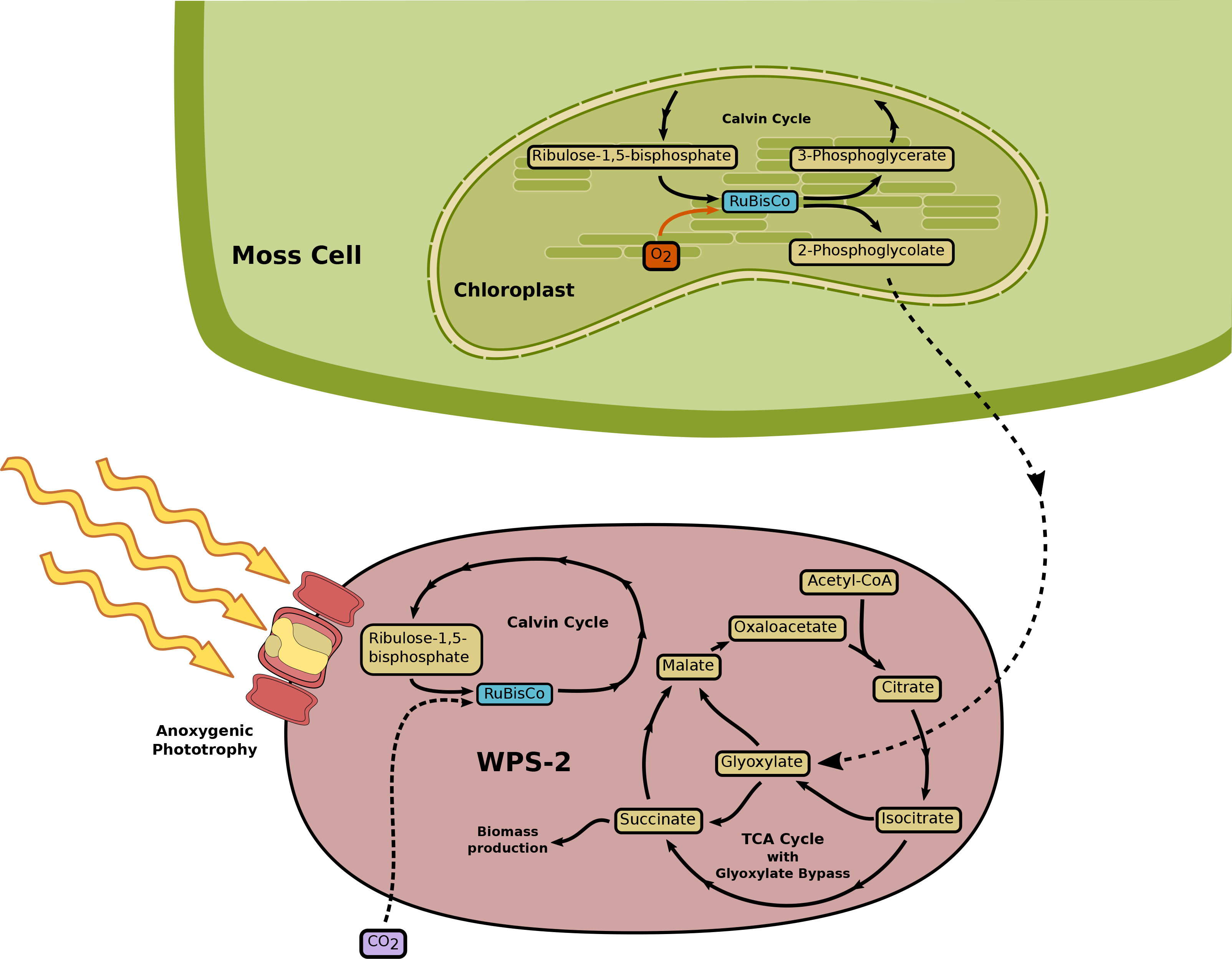
Diagram highlighting the inferred metabolic pathways linked to anoxygenic phototrophy and carbon cycling between the WPS-2 bacterial phylotype found to be abundant in mosses and the moss host cells. In the bacterial cell, light in a complementary spectrum to that absorbed by plants can be used to fuel cellular processes through anoxygenic phototrophy. CO_2_ fixation via RuBisCo in the bacterial cell may also occur, a novel feature for an aerobic anoxygenic phototroph. In the host cell, photorespiration (the action of RuBisCo on O_2_ rather than CO_2_) can produce a Calvin Cycle-inhibiting byproduct, 2-phosphoglycolate, that the bacterial cell may have the ability to metabolize for biomass production though conversion to glyoxylate, a process known as the “glyoxylate bypass”. This process represents a possible mechanism for a mutalistic or commensal interactions between these WPS-2 bacteria and the moss host.

If WPS-2 is an AAP possessing RuBisCo, how might it live in close association to moss and their cyanobacteria without being out-competed for light and carbon resources? WPS-2 appears to possess several traits which may allow it to effectively associate with mosses. As an anoxygenic phototroph, WPS-2 would not compete with its host for light resources since anoxygenic phototrophs absorb light in a complementary spectrum to that of plants and cyanobacteria, with a maximal absorption peak between 500 and 550nm (the region where chlorophyll has its minimum absorption) (Atamna-Ismaeel *et al.*, 2012). Interestingly, an absorption peak in this area of the spectrum may also give WPS-2 a photosynthetic advantage under light snow cover since the wavelengths with the greatest transmittance through snow are also in that region of the light spectrum (Perovich, 2007).

In addition to its complementary absorption spectrum, WPS-2 may be able to take advantage of the byproducts of its host’s photorespiration to supplement its C requirements. Photorespiration takes place when RuBisCo acts on O^2^ rather than CO^2^ producing phosphoglycolate, a potent inhibitor of the Calvin cycle. Thus, in plants and other carbon fixers who use the Calvin cycle, metabolizing phosphoglycolate is important. The moss-associated WPS-2 phylotypes appear to possess the enzymes necessary for converting phosphoglycolate into glyoxylate and passing it into the TCA cycle. This process, known as the “glyoxylate bypass”, is a modified TCA cycle that allows organisms to bypass the normal electron-generating steps of the cycle and devote it entirely to biomass production (Kornberg, 1966). Since WPS-2 is likely phototrophic, during times of high light it is unlikely to need its TCA cycle for energy generation and could instead take advantage of its host’s sugars for biomass production. In such a situation, WPS-2 would only need to use RuBisCo under conditions when sugars from the plants were scarce, such as early in the growing season or under snow pack. The complementary absorption spectrum of the moss-associated members of the WPS-2 phylum and their ability to metabolize glyoxylate appear to make this group uniquely equipped for living commensally on their moss hosts.

## Conclusions

We found that moss-associated bacterial communities are strongly structured by moss species identity with different moss species harboring distinct bacterial communities regardless of collection site. Surprisingly, all of the moss species studied possessed microbial communities that were capable of N_2_-fixation and hosted a broad diversity of putative N_2_-fixing bacterial lineages. However, it remains unclear which lineages are responsible for the measured N_2_-fixation rates and whether distinct bacterial lineages are driving N_2_-fixation in different moss species. Future work should focus on how microbial community variation between host species may influence N_2_-fixation rates and how interactions both between bacteria and between bacteria and the moss host might contribute to these rates. We also found that mosses can harbor a number of poorly-described bacterial lineages, including a high relative abundance of bacteria assigned to the candidate phylum WPS-2 with previously unknown ecological and metabolic characteristics. Using shotgun metagenomic analyses, we were able to assemble a nearly complete genome representative of this WPS-2 lineage and the genomic analyses suggest that WPS-2 is an anoxygenic phototroph that is uniquely adapted to living in close-association with mosses in this ecosystem.

## Acknowledgements

Financial support for this work was provided by the U.S. National Science Foundation Dimensions of Biodiversity program (DEB 1542609). The authors acknowledge infrastructural support provided by the University of Colorado Next Generation Sequencing Facility and the University of Alaska, Fairbanks. Helpful feedback on the manuscript was provided by the University of Colorado, EBIO writing co-op.

## Author Contributions

The project was conceived by N.F., M.C.M, and S.F.M. The manuscript was written by H.H.M. and N.F. with contributions from all co-authors. M.C.M. collected the samples. J.S. and S.M. performed the isotopic analysis. L.R.L. and S.F.M. identified the moss species and prepared the voucher specimens. H.H.M. and N.F. performed the molecular analyses.

## Conflict of Interest Statement

The authors declare no conflict of interest in this study.

## Materials and Correspondence

Requests for materials and correspondence should be addressed to N.F. or H.H.M.

## References

Atamna-Ismaeel, N., Finkel, O., Glaser, F., von Mering, C., Vorholt, J.A., Koblížek, M., et al. (2012) Bacterial anoxygenic photosynthesis on plant leaf surfaces. Environ. Microbiol. Rep. 4: 209–216.

Atamna-Ismaeel, N., Finkel, O.M., Glaser, F., Sharon, I., Schneider, R., Post, A.F., et al. (2013) Microbial rhodopsins on leaf surfaces of terrestrial plants. Environ. Microbiol. 14: 140–146.

Bay, G., Nahar, N., Oubre, M., Whitehouse, M.J., Wardle, D.A., Zackrisson, O., et al. (2013) Boreal feather mosses secrete chemical signals to gain nitrogen. New Phytol. 200: 54–60.

Baym, M., Kryazhimskiy, S., Lieberman, T.D., Chung, H., Desai, M.M., and Kishony, R.K. (2015) Inexpensive multiplexed library preparation for megabase-sized genomes. PLoS One 10: 1–15.

Bengtsson-Palme, J., Hartmann, M., Eriksson, K.M., Pal, C., Thorell, K., Larsson, D.G.J., and Nilsson, R.H. (2015) metaxa2: Improved identification and taxonomic classification of small and large subunit rRNA in metagenomic data. Mol. Ecol. Resour. 15: 1403–1414.

Bragina, A., Berg, C., and Berg, G. (2015) The core microbiome bonds the Alpine bog vegetation to a transkingdom metacommunity. Mol. Ecol. 24: 4795–4807.

Bragina, A., Berg, C., Cardinale, M., Shcherbakov, A., Chebotar, V., and Berg, G. (2012) *Sphagnum* mosses harbour highly specific bacterial diversity during their whole lifecycle. ISME J. 6: 802–13.

Bragina, A., Maier, S., Berg, C., Müller, H., Chobot, V., Hadacek, F., and Berg, G. (2012) Similar diversity of Alphaproteobacteria and nitrogenase gene amplicons on two related *Sphagnum* mosses. Front. Microbiol. 2: 1–10.

Brewer, T., Handley, K., Carini, P., Gibert, J., and Fierer, N. (2016) Genome reduction in an abundant and ubiquitous soil bacterium “*Candidatus Udaeobacter copiosus*.” Nat. Microbiol. 2: 16198.

Buckley, D.H., Huangyutitham, V., Hsu, S.F., and Nelson, T.A. (2007) Stable isotope probing with ^15^N2 reveals novel noncultivated diazotrophs in soil. Appl. Environ. Microbiol. 73: 3196–3204.

Caporaso, J.G., Lauber, C.L., Walters, W.A., Berg-Lyons, D., Huntley, J., Fierer, N., et al. (2012) Ultra-high-throughput microbial community analysis on the Illumina HiSeq and MiSeq platforms. ISME J. 6: 1621–1624.

Cole, J.R., Wang, Q., Fish, J.A., Chai, B., McGarrell, D.M., Sun, Y., et al. (2014) Ribosomal Database Project: data and tools for high throughput rRNA analysis. Nucleic Acids Res. 42: 633–642.

Dedysh, S.N. (2011) Cultivating uncultured bacteria from northern wetlands: Knowledge gained and remaining gaps. Front. Microbiol. 2: 1–15.

DeLuca, T.H., Zackrisson, O., Gentili, F., Sellstedt, A., and Nilsson, M.C. (2007) Ecosystem controls on nitrogen fixation in boreal feather moss communities. Oecologia 152: 121–130.

DeLuca, T.H., Zackrisson, O., Nilsson, M.-C., and Sellstedt, A. (2002) Quantifying nitrogen-fixation in feather moss carpets of boreal forests. Nature 419: 917–920.

Edgar, R.C. (2004) MUSCLE: Multiple sequence alignment with high accuracy and high throughput. Nucleic Acids Res. 32: 1792–1797.

Edgar, R.C. (2010) Search and clustering orders of magnitude faster than BLAST. Bioinformatics 26: 2460–2461.

Edgar, R.C. (2013) UPARSE: highly accurate OTU sequences from microbial amplicon reads. Nat. Methods 10: 996–8.

Geffert, J.L., Frahm, J.P., Barthlott, W., and Mutke, J. (2013) Global moss diversity: spatial and taxonomic patterns of species richness. J. Bryol. 35: 1–11.

Graham, E., Heidelberg, J.F., and Tully, B. (2017) Undocumented Potential For Primary Productivity In A Globally-Distributed Bacterial Photoautotroph. bioRxiv.

Grasby, S.E., Richards, B.C., Sharp, C.E., Brady, A.L., Jones, G.M., Dunfield, P.F., and Williamson, M.-C. (2013) The Paint Pots, Kootenay National Park, Canada — a natural acid spring analogue for Mars 1, 2. Can. J. Earth Sci. 50: 94–108.

Gundale, M.J., Deluca, T.H., and Nordin, A. (2011) Bryophytes attenuate anthropogenic nitrogen inputs in boreal forests. Glob. Chang. Biol. 17: 2743–2753.

Gundale, M.J., Nilsson, M., Bansal, S., and Jäderlund, A. (2012) The interactive effects of temperature and light on biological nitrogen fixation in boreal forests. New Phytol. 194: 453–463.

Ho, A. and Bodelier, P.L.E. (2015) Diazotrophic methanotrophs in peatlands: the missing link? Plant Soil 419–423.

Hughes, E., Head, B., Maltman, C., Piercey-Normore, M., and Yurkov, V. (2017) Aerobic anoxygenic phototrophs in gold mine tailings in. *Can*. J. Microbiol. 63: 212–218.

Ininbergs, K., Bay, G., Rasmussen, U., Wardle, D.A., and Nilsson, M.C. (2011) Composition and diversity of *nifH* genes of nitrogen-fixing cyanobacteria associated with boreal forest feather mosses. New Phytol. 192: 507–517.

Jean, M. (2017) Broadleaf litter and environmental effects on bryophytes in boreal forests of Alaska.

Jehmlich, N., Schmidt, F., Taubert, M., Seifert, J., Bastida, F., von Bergen, M., et al. (2010) Protein-based stable isotope probing. Nat. Protoc. 5: 1957–1966.

Joshi, N.A. and Fass, J.N. (2011) Sickle: A sliding-window, adaptive, quality-based trimming tool for FastQ files (Version 1.33) [Software].

Kip, N., Winden, J.F. Van, Pan, Y., Bodrossy, L., Reichart, G., Smolders, A.J.P., et al. (2010) Global prevalence of methane oxidation by symbiotic bacteria in peat-moss ecosystems. Nat. Geosci. 3: 617–621.

Kornberg, H.L. (1966) The role and control of the glyoxylate cycle in *Escherichia coli*. Biochem. J. 99: 1–11.

Kostka, J.E., Weston, D.J., Glass, J.B., Lilleskov, E.A., Shaw, A.J., and Turetsky, M.R. (2016) Tansley insight The *Sphagnum* microbiome: new insights from an ancient plant lineage. New Phytol.

Laforest-Lapointe, I., Messier, C., and Kembel, S.W. (2016) Tree phyllosphere bacterial communities: exploring the magnitude of intra- and inter-individual variation among host species. 4: 1–18.

Langmead, B. and Salzberg, S.L. (2012) Fast gapped-read alignment with Bowtie 2. Nat. Methods 9: 357–359.

Leff, J.W., Del Tredici, P., Friedman, W.E., and Fierer, N. (2015) Spatial structuring of bacterial communities within individual *Ginkgo biloba* trees. Environ. Microbiol. 17: 2352–2361.

Leppänen, S.M., Rissanen, A.J., and Tiirola, M. (2015) Nitrogen fixation in *Sphagnum* mosses is affected by moss species and water table level. Plant Soil 389: 185–196.

Li, D., Liu, C.M., Luo, R., Sadakane, K., and Lam, T.W. (2014) MEGAHIT: An ultra-fast single-node solution for large and complex metagenomics assembly via succinct de Bruijn graph. Bioinformatics 31: 1674–1676.

Lindo, Z. and Gonzalez, A. (2010) The Bryosphere: An Integral and Influential Component of the Earth’s Biosphere. 612–627.

Lundberg, D.S., Yourstone, S., Mieczkowski, P., Jones, C.D., and Dangl, J.L. (2013) Practical innovations for high-throughput amplicon sequencing. Nat. Methods 10: 999–1002.

Markowitz, V.M., Chen, I.M.A., Chu, K., Szeto, E., Palaniappan, K., Pillay, M., et al. (2014) IMG/M 4 version of the integrated metagenome comparative analysis system. Nucleic Acids Res. 42: 568–573.

Martin, M. (2011) Cutadapt removes adapter sequences from high-throughput sequencing reads. EMBnet.journal 17: 10–12.

McDonald, D., Price, M.N., Goodrich, J., Nawrocki, E.P., DeSantis, T.Z., Probst, A., et al. (2012) An improved Greengenes taxonomy with explicit ranks for ecological and evolutionary analyses of bacteria and archaea. ISME J. 6: 610–618.

Miller, C.S., Baker, B.J., Thomas, B.C., Singer, S.W., and Banfield, J.F. (2011) EMIRGE: reconstruction of full-length ribosomal genes from microbial community short read sequencing data. Genome Biol. 12: R44.

Op den Camp, H.J.M., Islam, T., Stott, M.B., Harhangi, H.R., Hynes, A., Schouten, S., et al. (2009) Environmental, genomic and taxonomic perspectives on methanotrophic Verrucomicrobia. Environ. Microbiol. Rep. 1: 293–306.

Opelt, K., Chobot, V., Hadacek, F., Schönmann, S., Eberl, L., and Berg, G. (2007) Investigations of the structure and function of bacterial communities associated with *Sphagnum* mosses. Environ. Microbiol. 9: 2795–2809.

Parks, D.H., Imelfort, M., Skennerton, C.T., Hugenholtz, P., and Tyson, G.W. (2015) CheckM: assessing the quality of microbial genomes recovered from isolates, single cells, and metagenomes. Genome Res. 25: 1043–55.

Perovich, D.K. (2007) Light reflection and transmission by a temperate snow cover. J. Glaciol. 53: 201–210.

Price, M.N., Dehal, P.S., and Arkin, A.P. (2010) FastTree 2 - Approximately maximum-likelihood trees for large alignments. PLoS One 5:.

R Core Team (2017) R: A Language and Environment for Statistical Computing.

Redford, A.J., Bowers, R.M., Knight, R., Linhart, Y., and Fierer, N. (2010) The ecology of the phyllosphere: geographic and phylogenetic variability in the distribution of bacteria on tree leaves. Environ. Microbiol. 12: 2885–93.

Rousk, K., Jones, D.L., and Deluca, T.H. (2013) Moss-cyanobacteria associations as biogenic sources of nitrogen in boreal forest ecosystems. 4: 1–10.

Rousk, K., Rousk, J., Jones, D.L., Zackrisson, O., and DeLuca, T.H. (2013) Feather moss nitrogen acquisition across natural fertility gradients in boreal forests. Soil Biol. Biochem. 61: 86–95.

Rousk, K., Sorensen, P.L., Lett, S., and Michelsen, A. (2015) Across-habitat comparison of diazotroph activity in the subarctic. Microb. Ecol. 69: 778–787.

Ruess, R.W., Anderson, M.D., Mcfarland, J.M., Kielland, K., Olson, K., and Taylor, D.L. (2013) Ecosystem-level consequences of symbiont partnerships in an N-fixing shrub from interior Alaskan floodplains. Ecol. Monogr. 83: 177–194.

Ruess, R.W., Mcfarland, J.M., Trummer, L.M., and Rohrs-Richey, J.K. (2009) Disease-mediated declines in N-fixation inputs by *Alnus tenuifolia* to early-successional floodplains in interior and south-central Alaska. Ecosystems 12: 489–502.

Sharp, C.E., Smirnova, A. V., Graham, J.M., Stott, M.B., Khadka, R., Moore, T.R., et al. (2014) Distribution and diversity of Verrucomicrobia methanotrophs in geothermal and acidic environments. Environ. Microbiol. 16: 1867–1878.

Trexler, R., Solomon, C., Brislawn, C.J., Wright, J.R., Rosenberger, A., Keddache, M., et al. (2014) Assessing impacts of unconventional natural gas extraction on microbial communities in headwater stream ecosystems in Northwestern Pennsylvania. Front. Microbiol. 5: 1–13.

Turetsky, M.R., Bond-Lamberty, B., Euskirchen, E., Talbot, J., Frolking, S., McGuire, a. D., and Tuittila, E.S. (2012) The resilience and functional role of moss in boreal and arctic ecosystems. New Phytol. 196: 49–67.

Vile, M.A., Kelman Wieder, R., Živković, T., Scott, K.D., Vitt, D.H., Hartsock, J.A., et al. (2014) N2- fixation by methanotrophs sustains carbon and nitrogen accumulation in pristine peatlands. Biogeochemistry 121: 317–328.

Wang, Q., Garrity, G.M., Tiedje, J.M., and Cole, J.R. (2007) Naïve Bayesian classifier for rapid assignment of rRNA sequences into the new bacterial taxonomy. Appl. Environ. Microbiol. 73: 5261–5267.

Wu, Y.-W., Tang, Y.-H., Tringe, S.G., Simmons, B. a, and Singer, S.W. (2014) MaxBin: an automated binning method to recover individual genomes from metagenomes using an expectation- maximization algorithm. Microbiome 2: 26.

Yurkov, V. and Hughes, E. (2017) Aerobic Anoxygenic Phototrophs: Four Decades of Mystery. In, Hallenbeck,P.C. (ed), Modern Topics in the Phototrophic Prokaryotes: Environmental and Applied Aspects. Springer International Publishing, Cham, pp. 193–214.

Yutin, N., Suzuki, M.T., and Béjà, O. (2005) Novel primers reveal wider diversity among marine aerobic anoxygenic phototrophs. Appl. Environ. Microbiol. 71: 8958–8962.

Yutin, N., Suzuki, M.T., Rosenberg, M., Rotem, D., Madigan, M.T., Süling, J., et al. (2009) BchY-based degenerate primers target all types of anoxygenic photosynthetic bacteria in a single PCR. Appl. Environ. Microbiol. 75: 7556–7559.

Zackrisson, A.O., Deluca, T.H., Nilsson, M., Sellstedt, A., Berglund, L.M., and Zackrisson, O. (2004) Nitrogen fixation increases with successional age in boreal forests. Ecology 85: 3327–3334.

Zackrisson, O., Deluca, T.H., Gentili, F., Sellstedt, A., and Jäderlund, A. (2009) Nitrogen fixation in mixed *Hylocomium splendens* moss communities. Oecologia 160: 309–319.

